# Opsin-free optical neuromodulation and electrophysiology enabled by a soft monolithic infrared multifunctional neural interface

**DOI:** 10.1101/2022.05.23.493057

**Authors:** Marcello Meneghetti, Jaspreet Kaur, Kunyang Sui, Jakob F. Sørensen, Rune W. Berg, Christos Markos

## Abstract

Controlling neuronal activity with high spatial resolution using multifunctional and minimally invasive neural interfaces constitutes an important step towards developments in neuroscience and novel treatments for brain diseases. While infrared neuromodulation is an emerging technology for controlling the neuronal circuitry, it lacks soft implantable monolithic interfaces capable of simultaneously delivering light and recording electrical signals from the brain while being mechanically brain-compatible. Here, we have developed a soft fibre-based device based on high-performance thermoplastics which are >100-fold softer than silica glass. The presented fibre-implant is capable of safely neuromodulating the brain activity in localized cortical domains by delivering infrared laser pulses in the *2 μm* spectral region while recording electrophysiological signals. Action and local field potentials were recorded *in vivo* in adult rats while immunohistochemical analysis of the tissue indicated limited microglia and monocytes response introduced by the fibre and the infrared pulses. We expect our devices to further enhance infrared neuromodulation as a versatile approach for fundamental research and clinically translatable therapeutic interventions.

## Introduction

Understanding the biological mechanisms underlying neural circuits is one of the fundamental goals in modern neuroscience [1]. Precise and reliable tools to modulate and monitor neuronal activities are essential elements within this frame [2, 3]. With the discovery of optogenetics [4, 5] and infrared neuromodulation (INM) [6, 7], light-induced control of neurons with high spatiotemporal resolution became one of the most powerful tools in neuroscience [8]. Therefore, new milestones have been set within the biotechnology community for novel devices able to deliver light in localized brain regions across the full electromagnetic spectrum [9-11]. Interrogation of the brain signals is a complex task and thus it requires bi-directional interfaces integrated with multiple functionalities in a single monolithic structure. An additional critical requirement for chronic *in vivo* experimentation is the suppression of the glial scarring by reducing the tissue’s foreign body response (FBR) [3]. Therefore, the design and development of minimally invasive devices based on biomaterials close to the Young’s modulus of the brain is crucial. [2, 12-14].

While optogenetics is arguably one of the most powerful methods for high-resolution modulation of neural populations, the necessity for genetic manipulation of the targeted cells adds further complexity to the process by requiring either the injection of a viral vector or the use of genetically modified animals [8, 15]. INM has thus attracted significant attention as a viable *transgene-free* alternative to optogenetics. This method relies on the light absorption from the biological tissue at specific water-dominant regions affecting the neuronal activity through temperature-mediated processes [16]. Moreover, INM offers tunable spatial resolution (by varying the operational wavelength) with the ability to affect individual axons without the need for the introduction of exogenous substances in the tissue [17, 18]. While the fundamental biophysical mechanisms of INM are not yet completely understood [7, 19], it is employed successfully in several biomedical research areas, such as restoration of hearing, investigation of brain diseases, control of peripheral nerves and cardiac pacing [7]. In the past few years, the aforementioned technological requirements for brain-compatible tools have been (to a large extent) addressed for use in optogenetics. Several groups around the globe have already reported innovative multifunctional brain devices such as polymer fibres and new soft planar implantable technologies. [3, 20-24]. However, developing of infrared (IR) counterparts to these technologies suitable for *in vivo* INM in the brain remains a challenge. IR light is normally delivered to regions deeper than the superficial cortical layers by implanting conventional silica optical fibres [24, 25] or silicon-based devices [26, 27]. Both platforms exhibit a significant mismatch in mechanical properties with respect to the brain tissue [2, 10]. Conventional step-index polymer optical fibres have been used in optogenetics and have in general found a vast amount of sensing applications in the visible regime [28]. However, this type of fibres suffers from significantly high absorption losses at longer wavelengths attributed to their fundamental vibrational absorption, which is inherently dependent on the chemical structure of the monomer. Consequently, the existing polymer optical fibre technology cannot be adopted for INM applications.

Here we address the abovementioned challenges of INM by developing a soft biocompatible multifunctional neural interface able to deliver IR light and record the brain activity *in vivo*, by thermally drawing a structured polymer optical fibre (SPOF) using non-conventional high-performance polymers. The developed SPOF devices can deliver light over a broad IR transmission spectral window spanning from 450 nm up to 2100 nm and overlapping with one of the main absorption peaks of biological tissues at 1930 nm. Microelectrodes have been integrated proximal to the fibre’s core, allowing *in situ* artifact-free electrophysiological recordings from the targeted region. Our SPOF implants delivered IR pulses from a supercontinuum laser, activating the neuronal circuitry with minimal tissue damage. The stability performance of the integrated microelectrodes during recordings of local field potentials (LFPs) was evaluated over multiple weeks of implantation.

## Results

### Fibre design and thermal drawing process

The choice of thermoplastic polymers suitable for thermal drawing needs to fulfill several optical, thermal and mechanical requirements such as refractive index difference to satisfy Snell’s law, biocompatibility, glass transition temperatures, and overlapping viscosity profiles for simultaneous thermal processing [29, 30].

The commonly used polymers in step-index optical fibres are polymethyl methacrylate (PMMA), polycarbonate (PC) and cyclic olefin copolymers (COCs) such as Topas and Zeonex [31]. Despite their relatively low optical loss in visible wavelengths (∼3.5 dB/m), they have never been reported to transmit light in the 2 μm range over a few centimeters, which are the typical required implantation length [14]. To fulfill the IR transmittance and flexibility requirements, we chose polysulfone (PSU) and fluorinated ethylene propylene (FEP) copolymer as materials for the fibre core and cladding, respectively. PSU is a sulfur-based polymer that provides good transmission in the IR compared to its more conventional optical polymer counterparts [14]. This high-performance polymer is characterized by high hydrolytic resistance and has been demonstrated to be highly biocompatible [32]. FEP, on the other side, is one of the softest biocompatible thermoplastics with Young’s moduli as low as 0.48 GPa, making it a promising material for suppressing the FBR while being optically transparent [33, 34]. Additionally, the large refractive index contrast between the two polymers leads to a very large numerical aperture (NA) (Fig. S1A), which is known to have an important role during neuromodulation [35]. The PSU and FEP viscosity-temperature profiles were characterized using standard rheometry and found to overlap and be suitable for fibre co-drawing (Fig. S1B).

The macroscopic template of the fibre (known as *preform*) was prepared by machining commercially available bulk polymer rods. Two side channels next to fibre’s core were introduced for the subsequent integration of the electrodes. The preform was then heated and scaled down into a fibre using an in-house draw tower facility (Fig. 1A). During the fibre fabrication, the drawing parameters of feeding/pulling speed were adjusted to obtain a core size of 105 µm, in order for the produced interfaces to be compatible with standard patch cables and components such as cannulae and adaptors typically used in animal experiments.

**Fig. 1.**
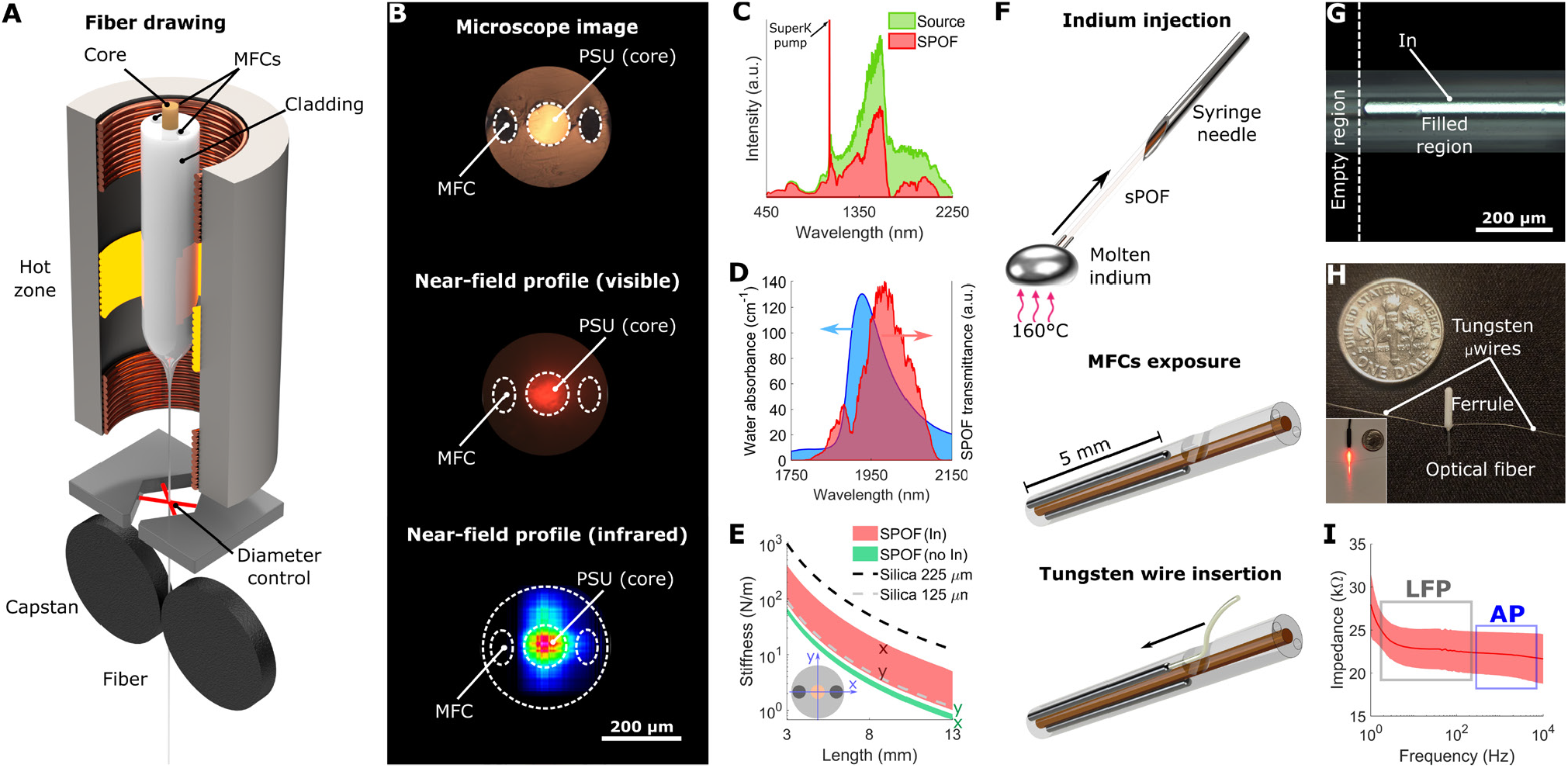
Fabrication and characterization of the multifunctional neural interfaces. (**A**) Schematic of the drawing process for the polymer optical fibre used to develop of the neural interfaces. (**B**) From top to bottom: microscope image of the structure of the fabricated optical fibre and near-field profiles of the output light at both visible and IR wavelengths. (**C**) Transmittance spectrum of the SPOF in the spectral range spanning from 450 to 2250 nm wavelengths. (**D**) Overlap between the absorbance of water (extracted from [55]) and the spectrum transmitted from the SPOF. The same light source and optical setup were used in the INS experiments shown later in the article. (**E**) Simulation of the bending stiffness of the SPOF, both with and without microfluidic channels filled with indium and compared with the most commonly used silica optical fibres in neuroscience. (**F**) Schematics of the fabrication steps to integrate electrodes in the optical fibre and connectorize them for EE recording. (**G**) Microscope image of the interface between the indium-filled and the hollow part of a microfluidic channel exposed for connectorization. (**H**) Assembled multifunctional probe after connectorization. Inset: visible light transmitted through the connectorized probe. (**I**) Impedance spectrum and corresponding standard deviation (shaded) of the integrated electrodes in the frequency ranges of LFPs and APs.

A single preform was enough to produce several hundred meters of SPOF with uniform core/cladding dimensions along the entire length. Minor deformations observed in the microfluidic channels can be attributed to the capillary forces acting on the FEP cladding during drawing combined with the blade fingerprint introduced by the cleaving (top image, Fig. 1B). The near-field profiles of the propagation modes in the visible (central image Fig. 1B) and IR (2 μm) band (bottom image, Fig. 1B) demonstrate the excellent confinement of the light in the core, which is crucial for enhanced localization of INM. The transmittance of the fibre was measured by coupling a coherent broadband supercontinuum source, showing two broad transmission windows whose combination covers from 450 nm up to 2100 nm (Fig. 1C). The long-wavelength spectral region (1750-2100 nm) contains most of the wavelengths commonly used in INM. By filtering the source with a long-pass filter (LPF) with a cut-off at 1800 nm, the output spectrum from the SPOF overlaps with the main absorbance peak of water at ∼1930 nm, as shown in Fig. 1D, which is the INM band used in this work. The mechanical softness of this fibre with hollow channels was also numerically evaluated, as further detailed in the next section (green shaded area, Fig. 1E).

### Preparation of the multifunctional neural interfaces

Electrophysiology is a critical modality to directly relate the neuronal response under different neurostimulation methods. Low-impedance electrodes were integrated in the SPOF, to enable simultaneous INM at the 2 μm spectral region and high-quality extracellular electrophysiology (EE) recordings. Short SPOF segments corresponding to the implantation length were connected to a syringe needle, which was used to create negative pressure in the channels to infiltrate molten indium (top, Fig. 1F). This specific metal was selected for the integrated electrodes due to its relatively low melting point (∼155 °C) and low Young’s modulus compared to other metals. Moreover, indium has been recently reported to be a suitable material for EE electrodes [21]. The choice of high-performance polymers with high melting points (>200 °C) as starting materials was critical in achieving the desired outcome, since it allowed the use of any metal with a melting point lower than the fibre’s materials as electrodes. Therefore, the SPOF could sustain the contact with the molten metal without any damage. The hollow channels were uniformly filled with indium, without introducing any structural deformation during the infiltration (Fig. 1G). The fibres were then extracted from the syringe and placed under an optical microscope to mechanically expose the hollow channels from the side (Fig. 1F, middle) and allow the insertion of 50 µm thick tungsten wires (Fig. 1F, bottom). Once the tungsten wires were in contact with the indium electrodes, they were fixed in position with cyanoacrylate adhesive. The SPOF neural interfaces were finally packaged for the *in vivo* experiments in ceramic ferrules (1.25 mm diameter, 6.4 mm length) (Fig. 1H) to achieve intuitive back-end connectorization. The impedance of the electrodes was characterized in the 1-10000 Hz range using electrochemical impedance spectroscopy in a three-terminal configuration (N=6). Values <35 kΩ were recorded in both the local field potential region (<250 Hz, [36]) and the action potential (AP) region (300-7000 Hz [37]) (Fig. 1I, S2). These values are well below the 1 MΩ at 1 kHz maximum value recommended for EE [10].

The bending stiffness of fibres correlates with tissue damage upon implantation and fixation to the skull [14]. Therefore, its value was determined using a finite element method (FEM) model for different lengths based on cross-sectional geometry. These calculations were performed both for the as-drawn SPOF and for the implants with integrated metallic electrodes, by changing the mechanical parameters of the hollow channels between air and indium. Despite the presence of metallic electrodes and the large overall diameter of the fibre, the calculated stiffness was found to be comparable to or even lower than the one of the silica fibres with diameters most commonly used in optogenetics applications (125 and 225 µm external diameter) (Fig. 1E). Since, unlike silica fibres, the SPOF and implants are not centrosymmetric their stiffness is represented as a shaded area to account for its angular dependence. The lower edge of the area represents the minimum stiffness level, resulting from a bending force parallel to the axis connecting the center of the electrodes in the fibre’s cross section. In contrast, the upper edge represents the maximum, i.e., a bending force perpendicular to that same axis.

### In vivo *simultaneous INM and electrophysiology*

Water absorption is the main factor introducing a thermal response to IR light in biological tissues. Our neural interfaces were tested *in vivo* to evaluate their performance for modulation of the neural activity in the cortex of adult rats with IR light overlapping with one of the strongest water absorbance peaks in the 2 μm region (Fig. 1D). During modulation, the integrated electrodes recorded the EE response of neurons to the IR light (Fig. 2A) delivered >1mm deep into the brain following implantation of the interfaces in anesthetized animals during a terminal procedure (Fig. 2B). The INM protocol employed consisted of 2 minutes illumination periods, similar to [26], interspersed by 2-3 minutes recovery periods. This investigation was repeated at four distinct cortical locations, using four different average power levels (10 mW, 12.5 mW, 15 mW and 17.5 mW). Since the brain tissues had to be further analyzed by immunohistochemistry (IHC) to evaluate the laser-induced damage (as presented in the following section), no positional micro-adjustments were made once the implant was in the desired region. This way, the surgery-related damage that might have masked the laser-induced one was minimized. The suitability of our multifunctional neural interface to investigate neuronal dynamics in response to light-induced heating was verified by recording EE signals at the aforementioned four different INM powers (Fig. 2C). The spike activity was recorded consistently and repeatably for each power level over several stimulation cycles (Fig. S3). At the highest power of 17.5 mW, we recorded the potentials and identified single-unit neuronal activity (Fig. S4). In contrast, with 10 mW, 12.5 mW and 15 mW, the recorded signals have a less defined and more variable shape, compatible with multi-unit activity. The setting of the recording setup, targeted towards single unit APs, strongly affected the signal-to-noise ratio (SNR) for these recordings; therefore a digital low-pass frequency filter was applied to improve the visualization of the spikes at 10, 12.5 and 15 mW stimulation power in Fig. 2C. While effectively suppressing noise, this data post-processing introduced a reduction in the plotted intensity and a slight change in the spikes’ shape, though leaving the temporal position of the spikes unaltered (Supplementary text 1, Fig. S5 and S6). Furthermore, the presence of a stimulation-activity time delay, the shape of the signal, and recordings with no activity allow us to conclude that the recorded signal originated from the IR-modulated neural activity and not from any other side laser, electrical or thermal artifacts.

**Fig. 2.**
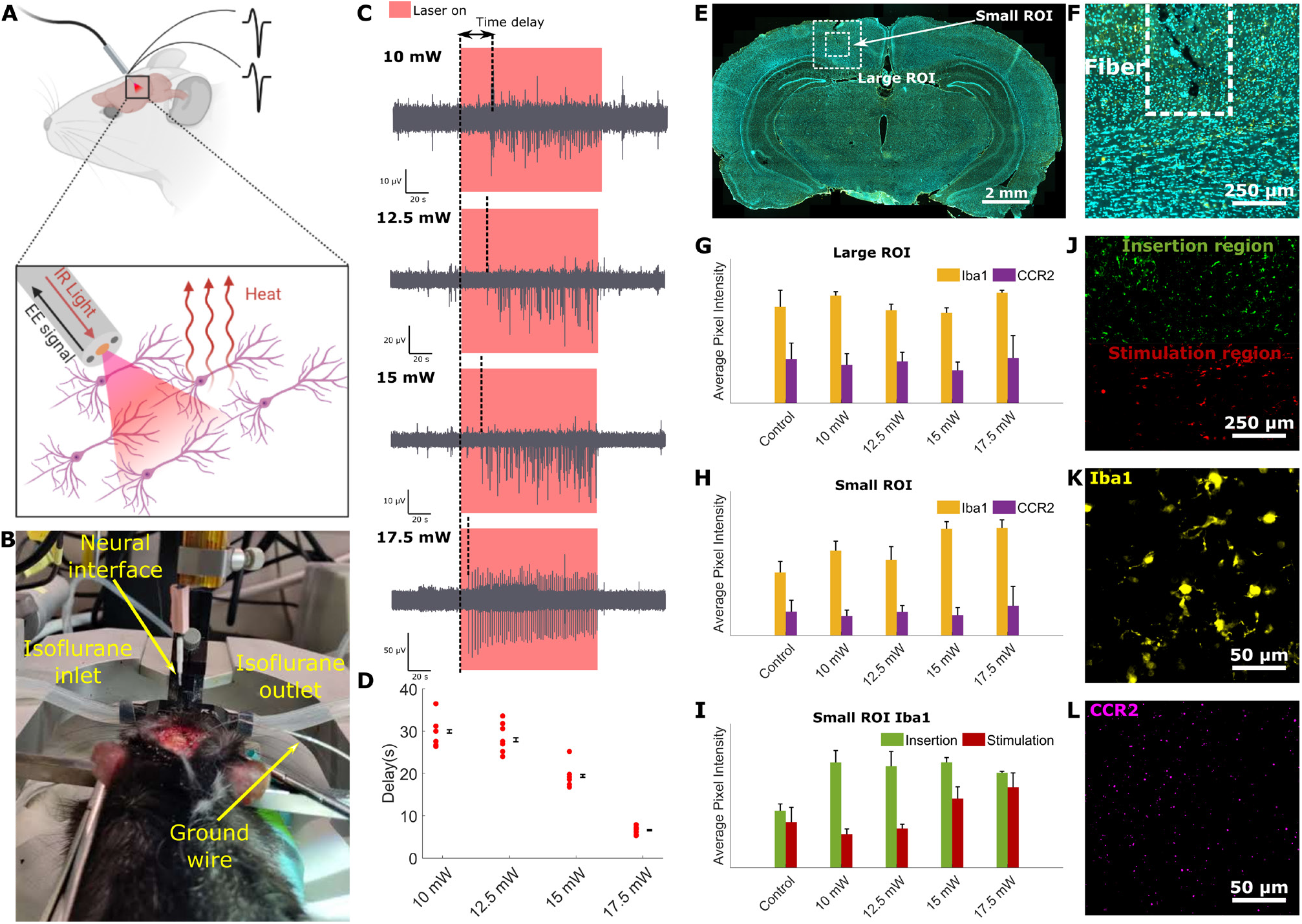
Opto-electrophysiological interrogation of the brain. (**A**) Concept of the experiment: delivering IR light with the developed interface to generate heat-induced variations in neural activity while conducting simultaneous electrophysiology based on the integrated electrodes. (**B**) Picture of the surgery for the insertion of the implant in the brain, taken immediately prior to the insertion. (**C**) Neural response to two minutes stimulation cycles for laser powers ranging from 10 to 17.5 mW. (**D**) Red dots: delays between the beginning of the stimulation and the beginning of the recording activity for all stimulation cycles at different powers. The average and standard error are presented on the side of each series. (**E**) Microscope image of a brain slice used for IHC, stained with Iba-1 (yellow) and DAPI (cyan), with the ROIs used for analysis indicated as dashed lines. (**F**) Magnification of 2F showing the small ROI only; the dashed lines indicate the position of the fibre during stimulation. (**G, H**) Average fluorescence intensities for Iba-1 and CCR2 in the large and small ROIs, respectively (N_slices_>5). (**I**) Average fluorescence intensities for Iba-1 in the insertion and stimulation regions of the small ROI (N_slices_>5). (**J**) Colorized depiction of a small ROI’s subdivision into insertion and stimulation regions (Iba-1 channel only). (**K, L**) Magnified images showing microglia and monocytes in the implantation region, respectively.

While neural responses to INM are often reported in the literature to be in the millisecond timescale [18, 38], we observed a seconds-long time delay between the light delivery and onset of the stimulated activity. The short (picosecond) laser pulses used in this investigation can explain the significantly large response time. This class of pulses is known to induce both thermal and stress confinement (Supplementary text 2), giving rise to complex light-matter interaction dynamics within the realm of photothermal and photomechanical effects that might delay the temperature increase. Furthermore, the response time decreases from ∼30 to 7 seconds as the output power increases from 10 to 17.5 mW (Fig. 2D). This effect was verified by repeatedly measuring the brain activity over several stimulation cycles (Fig. S3).

Since INM offers high spatial resolution for targeted stimulation, it was recently employed to map mesoscale brain connectomes using functional magnetic resonance imaging (fMRI) in anesthetized animal (cat) [19]. To investigate the time-resolved dynamics of the brain during the previously described seconds-long response time of the electrical activity, we also conducted fMRI during INM. The experiment was conducted in the motor cortex of an anesthetized rat to visualize the brain region, using an optical power of 15 mW for INM. The requirement of relatively short acquisition times (∼2 s) to temporally resolve the onset dynamics of the stimulation, negatively affected the maximum achievable spatial resolution (0.5×0.5×1 mm) of the obtained images. However, it was possible to verify the presence of blood-oxygen-level-dependent (BOLD) signal fluctuations temporally correlated with the illumination in the voxels close to the fibre tip (Fig. S7).

### Evaluation of laser-induced damage

Since the INM experiments performed in this investigation use a non-standard protocol with a supercontinuum laser (broadband, picosecond long pulses in the IR) delivered in the brain tissue, as opposed to the monochromatic lasers normally used in literature [18,19,26], it is important to investigate the neuronal damage after INM. This damage assessment was conducted by post-experimental transcardial perfusion of the rat, out-dissection of the central nervous system, slicing of the brain and IHC analysis. Due to the terminal nature of the experiment, the brain slices were stained with two widely used markers to identify acute inflammation: activated macrophage marker ionized calcium-binding adaptor molecule 1 (Iba-1) and monocyte marker C-C chemokine receptor 2 (CCR2). Pristine rat brains were similarly perfused, dissected and analyzed as a control. We subsequently used the fluorescence intensity in square regions surrounding the SPOF tip as an indicator to quantify the tissue damage. To visualize the effects of the procedure as a whole, in terms of both the insertion and the laser-induced thermal load, the average pixel intensity was firstly measured in a region of interest centered on the fibre’s tip and comprising the whole length of the implant (large region of interest, ROI) (Fig. 2E). However, a large portion of unaffected tissue was included in the analysis, leading to the lack of any statistically significant difference between the control brains and the brains that underwent surgery and INM (Fig. 2G). We therefore reduced the region of interest to 1×1 mm, while keeping it centered on the fibre’s tip (small ROI, Fig. 2E-F). In this smaller region, while the intensity related to CCR2 remained similar for all the sample groups, an increase in Iba1-related intensity was observed for the stimulated samples, showing the presence of acute inflammation induced by the procedure (Fig. 2H). Since the focus of this work is on the laser-induced damage, the small ROI was split perpendicularly to the fibre into an insertion region and a stimulation region (Fig. 2I-J), thus effectively separating the effects of the laser heating from the mechanical damage caused by the implantation surgery. A constant increase in Iba-1 intensity was observed in the insertion region for all the stimulated samples, compatibly with the fact that the insertion procedure was identical in all brain regions. In contrast, a power-dependent increase of the intensity was measured in the stimulation region, where the IR thermal load is expected to be directly proportional to the laser power. The fluorescence signals from the Iba-1 and CCR2 markers are shown in Fig. 2K and 2L, respectively.

### Measurement of local field potentials using the integrated electrodes

Given that the main aim of developing our multifunctional neural interfaces is to enable chronic INM studies, we investigated the long-term stability of the interface between the integrated electrodes and the brain. Since LFPs have attracted a strong interest in the last decade from the community for adaptive neuromodulation [39, 40], as a verification method we quantified the LFPs’ SNR over four weeks of recording in behaving animals. Hence, the neural interfaces were chronically implanted in the left hippocampus of 2 rats. Tungsten microwires (µwires), which are highly stable electrodes and commonly used in the literature for LFP recording [41-43], were also implanted in the right hippocampus to provide a benchmark (Fig. 3A). The flexible µwires (50 µm diameter) were placed in stainless steel cannulae similar in size to our interfaces, to ensure correct positioning during implantation (Fig. 3B). Representative 5 seconds periods were extracted from each recording, using a high-pass filter (1Hz) to remove baseline drifts and DC offsets (Fig. 3C). As shown in Fig. 3D-3G, the recorded oscillations had their main spectral components in the 6-10 Hz region, directly related to the emergence of hippocampal theta rhythm correlated to the animal’s movements [44, 45]. The SNR used as a stability indicator was defined as the ratio between the total collected signal falling in the 6-10 Hz window and the total signal outside these frequencies. The trend of the SNRs calculated from the 5-seconds periods over time show a similar performance between the integrated electrodes and the tungsten µwires in terms of stability (Fig. 3H). Furthermore, the brains were sliced and imaged at the end of the four weeks to provide a visual estimation of the scarring left in the brain by the two different kinds of chronic implants (Fig. S8).

**Fig. 3.**
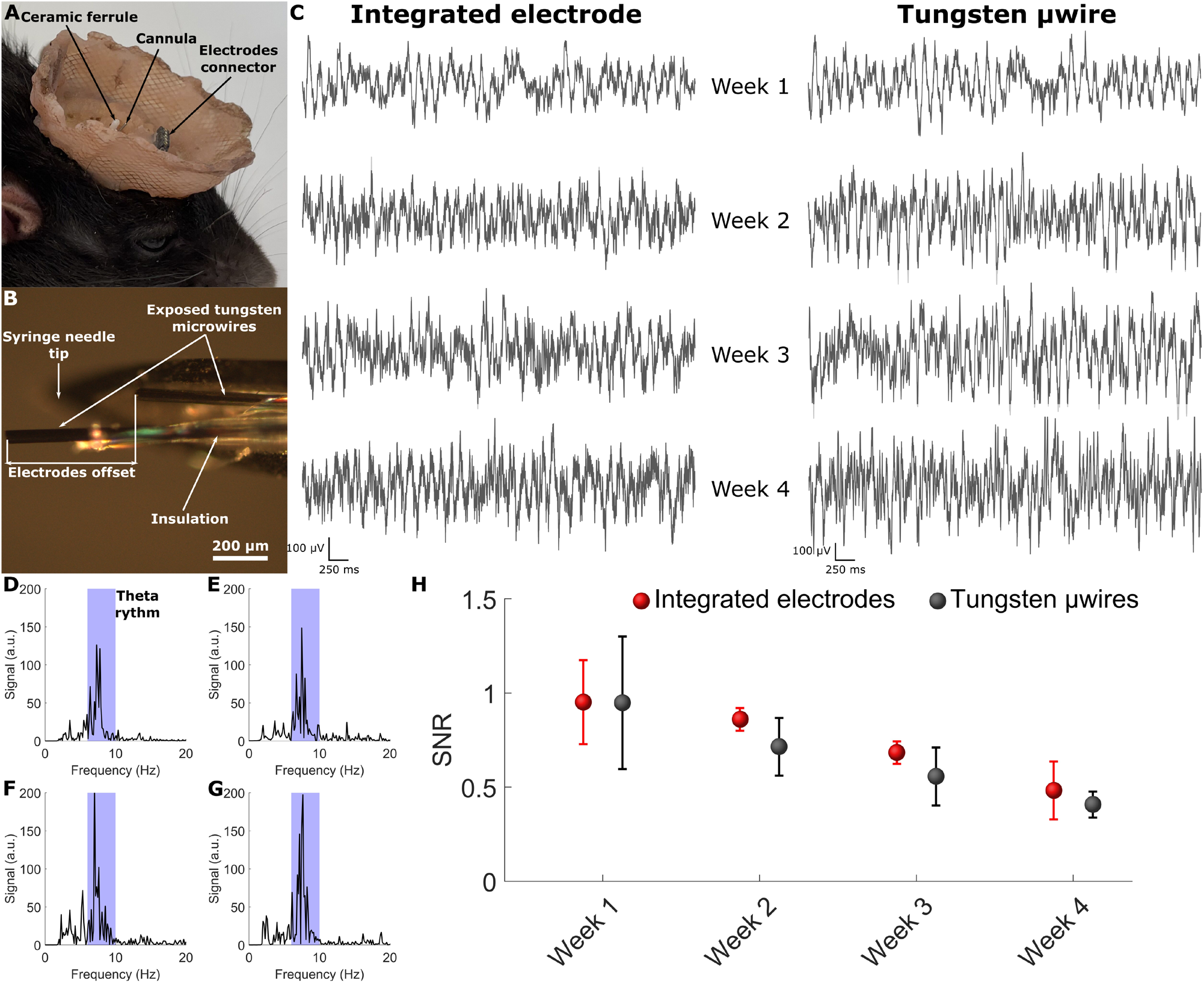
Chronic local field potential measurement in the hippocampus. (**A**) Chronic implant of a multifunctional neural interface and a steel cannula with tungsten µwires in the brain. (**B**) Magnified image of the tip of one of the cannulae used for this experiment. (**C**) 5-second-long EE recordings of LFPs in the hippocampus from an integrated indium electrode and a tungsten µwire over four weeks. (**D-G**) Fast Fourier transform of the indium electrode recordings shown in Fig. 3C for weeks 1, 2, 3 and 4 respectively, with the 6-10 Hz region corresponding to theta rhythm highlighted in light blue. (**H**) Comparison of the calculated signal-to-noise ratio over the four weeks for integrated electrodes and tungsten µwires (n=4 for both electrode types), with error bars showing the standard error.

## Discussion

We have developed SPOF-based IR multifunctional neural interfaces for INM applications for the first time. The initial fibre templates were fabricated by high-performance thermoplastics using a thermal drawing process. This method ensures the scalability, as well as the future possibility to easily reduce the device’s diameter to further suppress FBR. The latter claim is also supported by the fact that electrodes that were integrated in the fibre had a low measured impedance, leaving space for reduction of their surface without compromising the recording performance. Furthermore, the high thermal performance and hydrolytic stability of the polymers used to fabricate the SPOF make the device suitable for sterilization both by steam heat and dry heat [46, 47], which are between the most cost-effective and most diffused sterilization methods for medical devices [48], contributing thus to the efforts for sustainable biotechnology.

Our neural interfaces were verified by *in vivo* simultaneous INM and EE in anesthetized and behaving animals. A supercontinuum source covering a broad transmission spectrum was used to enable stimulation at the water absorption peak at the 2 µm region. This protocol has been here proven to efficiently and reproducibly evoke neural activity at different powers, with minimal power-dependent inflammation in the brain, confirming the existing literature on INM being a safe approach for neuromodulation under controlled conditions. The integrated electrodes allowed us to obtain EE recordings with low noise (<30 µV peak-to-peak). The use of IR light efficiently removed the artifacts commonly arising from the Becquerel effect when using shorter (visible) wavelengths [49].

The hollow channels in the SPOF were functionalized with metal electrodes to develop a protocol for the straightforward integration and back-end connectorization of the electrical recording functionality in our neural interfaces. It should be noted that the resulting indium-tungsten electrodes are highly reproducible as shown by the impedance spectra. Testing in freely behaving animals showed that both the electrode connections and the brain-electrode interfaces are as efficient and reliable as other solutions that are already commonly employed in the field of neuroscience.

INM is a powerful tool to investigate the function of small structures in the brain by affecting both excitatory and inhibitory circuits [19]. Our soft neural interfaces could greatly advance the use of this technique in deep brain regions and chronic settings, to gain a better understanding of neural mechanisms at a circuit level. Furthermore, due to the biocompatibility and sterilizability of the used materials, they may also increase the potential for translation of the findings towards the development of new therapies for neural diseases in humans.

## Methods

### Mechanical and optical characterization of the materials

The viscosity-temperature curves of PSU (core) and FEP (cladding) were measured in the 40-300°C temperature range using a rotational rheometer (TA, Discovery Hybrid HR-2). Thin rectangular samples of dimensions 2 mm thick, 10 mm wide and 25 mm in length were loaded into the clamps of the system and locked with screws. A constant upward force is applied in the axial direction to analyze the solid samples. We chose an axial force of 1 N with a sensitivity of 0.1 N. The refractive indices of the materials were measured in bulk hot-embossed samples in the 210-1000 nm range using an ellipsometer (J.A.Woollam, VASE) with a 5-nm resolution [50].

### Thermal drawing of the IR fibres

Commercially available rods of FEP and PSU were purchased by Goodfellow, UK and annealed in a drying oven for more than two weeks at 120 °C. The preform was assembled by machining the rods to obtain a 15 cm long FEP cylinder with a total diameter of 26.2 mm, a 7 mm central hole and two 6 mm side holes, and a solid PSU cylinder with a diameter of 7 mm. The fibre was thermally drawn using a three-zone furnace by stabilizing the temperature of the central one at 260°C. The preform was heated and pulled at a speed of 2 meters/minute. The fibre diameter was controlled during the process using a high-precision laser detector (< 0.1 µm resolution).

### Integration of the indium electrodes

The integration of the indium electrodes in the SPOF started by cutting 5-cm-long pieces of fibre and partially inserting them in 18G syringe needles. The fibre pieces were subsequently sealed in position using epoxy resin (G14250, Thorlabs), which was left to dry overnight. Afterwards, a short length of indium wire (diameter 1.5 mm, 99.999% purity, Goodfellow, UK) was melted on a heating plate and infiltrated into the side channels over >5 mm by using a syringe. The fibre pieces were then separated from the needles with a blade, and the filling quality was verified with an optical microscope, under which the non-filled length of the hollow channels was exposed on both sides by using a scalpel. Tungsten wires (diameter 50 µm, purity 99.999%, Goodfellow UK) were manually inserted until contact with the indium was achieved and were finally glued in position with small drops of cyanoacrylate adhesive (Loctite).

### Fabrication of the tungsten µwire cannulae

As a first step to fabricate the tungsten µwire cannulae used as a benchmark for LFPs recording, the coating was removed from a 300 µm length at one end of 10-cm-long pieces of insulated tungsten wires (A-M Systems, 50 µm bare diameter, 100 µm coated diameter). The wires were then inserted in a 26G syringe needle acting as a guide for the implantation. The wires were finally fixed with glue with a ∼500 µm offset between the wire tips, to avoid short-circuiting during the implantation.

### Microscope imaging of the fibre and near field profiles

To image the cross section of the fabricated fibres, we first cleaved short lengths of the fibres and ground them on different grits of polishing paper, starting from 30 μm down to 0.3 μm. The polished fibre cross-sections were then imaged using a Zeiss Axioscan A1 microscope, equipped with an Axiocam 305 color camera, in transmission mode. The image of the fibre’s side in Fig. 2G was acquired using the same microscope in reflection mode.

The near field profile of the propagating light in the SPOF was recorded in the visible by coupling light from a Thorlabs M660L4 red LED and imaging the collimated output light with an IDS U3-3680XLE camera. A similar procedure was used for the near IR near field profile, using a BKtel 1.55 µm custom nanosecond laser as a light source and recording the output with a Thorlabs BP109-IR2 slit scanning beam profiler.

### Broadband transmission measurements

The SPOF’s optical transmission was measured by butt-coupling the output light from a broadband supercontinuum source (NKT SuperK Extreme) in the SPOF’s core, using a silica patch cable with the samecore diameter of 105 µm (Thorlabs M61L01). Spectra at the SPOF’s output were recorded using an SP320 scanning spectrometer from Instrument Systems.

### Numerical modelling of bending stiffness

To evaluate the flexibility of the fabricated SPOF, a FEM analysis model based on COMSOL Multiphysics was used to simulate the bending and calculate the corresponding bending stiffness. 3D FEM models of the fibre, both with and without the indium electrodes, were developed starting from the cross-sectional geometry of the drawn fibre. For the modeling of standard commercial fibres, uniform silica cylindrical shapes were used as geometry, since there is almost no variation in mechanical properties between core and cladding. Similarly to the one used in [35], the numerical model is based on fixing one end of the fibre while applying a force perpendicular to the fibre’s axis to the other end. The simulated diameters were 400 µm for the SPOF and 125 and 250 µm for the silica optical fibres.

### Impedance spectroscopy

The impedance characterization of the integrated electrodes was conducted in phosphate-buffered saline (PBS) using a Hioki IM 3590 chemical impedance analyzer in a three-terminal configuration (Fig. S9). A tungsten microwire wrapped around a glass rod acted as a counter electrode and an Ag/AgCl electrode was used as a reference electrode. The frequency range used for the measurements was 1-10000 Hz.

### Animals and ethical statement

Wild-type adult Long Evans rats (Charles River Laboratories) were used to perform experiments. All the procedures concerning the housing of rats, habituation in the animal facility, surgery and post-surgery care were approved by the national veterinary and food administration (permission number for the animal research: 2019-15-0201-00018) and in compliance with the European Union Directive 2010/63/EU.

### Pre-implantation procedures

All surgical procedures were aseptic. Anesthesia was induced in the rats using isoflurane gas (1–3% in oxygen), followed by head shaving and skin disinfection with chlorohexidine followed by 70% ethanol. The animals were then moved to a stereotaxic frame with a heating pad and oximeter to maintain and monitor the animal. After confirming surgical anesthesia by loss of toe pinch response, the head was fixed using ear bars. Lidocaine was administered subcutaneously at the incision location, and ocryl gel was applied to maintain eye moisture. A sterile drape was placed to cover the entire body, followed by further cleaning of the head with 70% ethanol. Finally, an incision was made to expose the skull. Anesthesia was maintained throughout the surgery by constant administration of isoflurane gas at the abovementioned concentration.

### Acute implantations

For acute implantations (n=2), a surgical drill was used to expose four distinct regions of the brain cortex (locations shown in Fig. S10) and a posterior brain region for grounding the animal with a silver wire. A minimum of 4 mm distance between the individual regions was maintained. The multifunctional neural interface was inserted sequentially in the different regions at a depth of ∼1 mm, using a single smooth motion to avoid additional damage to the tissue, to perform the simultaneous INM and EE experiments at different powers. At the end of the experiments, the rats were deeply anesthetized by an intraperitoneal injection of pentobarbital, followed by transcardial perfusion [51, 52] with a solution of 4% paraformaldehyde (PFA) in PBS.

### Chronic implantations

For chronic implantations, 3% hydrogen peroxide was applied to the skull, clearing connective tissue, followed by thorough washing with saline solution. Electrocautery was used to stop any bleeding. A 3D printed baseplate was adhered to the skull using Metabond (Parkell) cement, and a fine sheet of copper mesh was glued onto the baseplate. Using the stereotaxic robot and drill, a small hippocampal area (2 mm diameter) was exposed on each hemisphere (anterioposterior 3.6 mm, mediolateral 2 mm, dorsoventral 3 mm from dura mater). A multifunctional neural interface was implanted in the left hemisphere while a cannula with tungsten µwires was implanted in the right hemisphere. The skull was grounded using a silver wire in contact with the cerebrospinal fluid. All the regions were sealed using silicon adhesive (Kwik-Sil, World precision instruments) followed by cyanoacrylate adhesive and dental cement. Finally, the copper mesh was molded into a crown, covered by dental cement to smoothen the surface (Fig. 3A). The crown served two functions: the protection implants from damage during the movement of the rat and the minimization of electromagnetic noise while recording electrical signal from the brain (by acting as a Faraday cage). After the surgery, the rats were orally treated with buprenorphine (0.2 mg mixed with 1 g Nutella) every 12 hours for three days. Furthermore, Carprofen (5 mg/kg), and Baytril (5 mg/kg) were administered subcutaneously every 24 hours for 5 and 10 days, respectively. The conditions of the animals were monitored 2-3 times per day over the first 3 days and once daily for the following week.

### In vivo IR neuromodulation

Light from a broadband supercontinuum source (NKT SuperK Extreme, 1-20 MHz repetition rate, average power up to 6W) was filtered using a long-pass filter (cut-off wavelength of 1800 nm) and coupled to a silica fibre patch cable. The patch cable was then connected to a multifunctional neural device using a ceramic mating sleeve. The power at the output of the devices was recorded prior to each insertion using a Thorlabs S401C thermal power sensor. Following insertion, light was delivered continuously for 2 minutes intervals, followed by 2-3 minutes resting periods. Four distinct regions of the brain cortex were stimulated at different optical powers (10, 12.5, 15 and 17.5 mW) for each animal, starting with the highest power and proceeding sequentially towards the lowest one.

### Electrophysiological recordings

The acute EE recordings during INM were conducted by connecting the exposed tungsten wires from the neural devices to the headstages of differential amplifiers (DP-311A, Warner Instruments). The signal was recorded through an ADInstruments PowerLab 4/26 DAQ using LabChart 8 software. For all recordings, the internal high-pass and low-pass filters of the amplifiers were set at 300 Hz and 10 kHz respectively, and the sampling rate was set at 20 kHz.

The chronic electrophysiological recordings of LFPs in freely behaving animals were performed using an Intan RHD 2000 USB interface board equipped with a RHD 2164 amplifier, at a sampling rate of 20 kHz. The data was recorded using Intan’s RHX Data Acquisition Software.

### Immunohistochemistry

After the acute experiments, the animals’ brains were removed and kept in PFA for 4 h before being transferred to sucrose 30% (w/v) for cryoprotection. A cryostat was used to slice coronal brain sections (25 μm thickness) that were immediately collected on superfrost plus glass slides (Thermo Fisher Scientific GmbH, Germany) for IHC following a procedure similar to [53, 54]. The slices were washed in PBS and incubated a room temperature (2 h) with a blocking solution (5% bovine serum albumin, 0.3% Triton X-100, 5% fetal bovine serum, 1% PBS). Anti-CCR2 (1:100, rabbit polyclonal, Thermo Fisher Scientific PA5-23037) and anti-Iba-1 (1:1000, rabbit polyclonal, Wako 019-19741) primary antibodies were applied to the slices, followed by overnight incubation at 4 °C. Afterwards, the slices were rewashed in PBS and secondary antibodies (donkey anti-rabbit, Alexa Fluor 633, 1:500, Invitrogen) were applied, together with 4’,6-Diamidino-2-Phenylindole, Dihydrochloride (DAPI, Thermo Fisher Scientific, D1306) was applied at a concentration of 1:1000. Finally, after 2 h at room temperature, the glass slides were mounted using DAKO mounting medium and imaged using a Zeiss Axioscan Z1 microscope (20X magnification).

The IHC images were initially analyzed using the Fiji open-source image processing package, based on ImageJ2. All images were opened at full resolution in the software, and the marker and DAPI channels were separated. The marker channel images were then cropped to two different square ROIs (referred to as “large” and “small” in the text) of 2×2 mm and 1×1 mm sizes, respectively, centered around the tip of the inserted SPOF. After discarding the images found to be out of focus or presenting too large artifacts, a threshold was applied to eliminate the signal from out-of-plane cells, and the maximum pixel value was adjusted to increase contrast. The threshold and maximum were kept as constants for all images obtained from slices stained with the same marker, while variating between different markers. The average pixel intensity was then calculated using Fiji. For a more precise analysis, the small ROIs for slices stained for the Iba-1 marker were further divided into 0.5×1 mm ROIs (insertion and stimulation), splitting them across a line normal to the fibre insertion direction.

### fMRI imaging and analysis

The fMRI data was acquired on a 9.4 T scanner (Bruker Biospec 9.4/30 USR AVANCEIII/MRI) during a stimulation protocol similar to the one described for simultaneous INM and electrophysiology (15 mW, 2 minutes OFF – 2 minutes ON cycles, 5 repetitions). Structural images were obtained using a TurboRARE sequence with an in-plane resolution of 0.137×0.137 mm (5 slices, 0.5 mm thickness), a repetition time of 3500 ms and an echo time of 34 ms. The bandwidth for these measurements was 50 kHz, and the matrix size was 256×256 (field of view 3.5×3.5 mm). Functional images were obtained with a spin echo EPI sequence at a voxel size of 0.5×0.5 (in plane) x 1 (slice thickness) mm (3 slices) in a 64×64 matrix. The echo and repetition time were of 35 ms and 2000 ms respectively, the bandwidth was 250 kHz, and the field of view was of 32×32 mm.

Functional MRI images were opened as stacks in the Fiji software, and the slices centered around the fibre insertion region were isolated. The temporal profiles of voxels in a 5×7 voxel region of interest close to the fibre tip were then exported from Fiji as numerical vectors for further analysis.

### Data processing and representation

The data processing, analysis and plotting were performed in the Matlab 2019b suite.

## Acknowledgments

This work was financially supported from Lundbeck Fonden projects (Multi-BRAIN, R276-2018-869) and from VILLUM FONDEN (36063). We thank Yuki Mori, Palle Koch and Ryszard S. Gomolka from Panum NMR Core Facility for their technical support. We also thank Guanghui Li from the Department of Neuroscience at Copenhagen University, as well as Ole Bang and Yazhou Wang from the Department of Electrical and Photonics Engineering at the Technical University of Denmark, for providing their expertise and support during the study.

## Data availability

The data supporting the results in this study are available within the paper and its Supplementary Information.

## Competing interests

The authors declare no competing interests.

## Notes

### Competing Interest Statement

The authors have declared no competing interest.

## References

1. Vázquez-Guardado, A., Yang, Y., Bandodkar, A. J. & Rogers, J. A. Recent advances in neurotechnologies with broad potential for neuroscience research. Nature Neuroscience 23, 522–1536 (2020).

2. Won, S.M., Song, E., Reeder, J. T. & Rogers, J. A. Emerging modalities and implantable technologies for neuromodulation. Cell 181, 115–135 (2020).

3. Sung, C., Jeon, W., Nam, K. S., Kim, Y., Butt, H. & Park, S. Multimaterial and multifunctional neural interfaces: from surface-type and implantable electrodes to fiber-based devices. Journal of Materials Chemistry B 8, 6624–6666 (2020).

4. Zemelman, B. V., Lee, G. A., Ng, M. & Miesenböck, G. Selective photostimulation of genetically chARGed neurons. Neuron 33, 15–22 (2002).

5. Boyden, E. S., Zhang, F., Bamberg, E., Nagel, G. & Deisseroth, K. Millisecond-timescale, genetically targeted optical control of neural activity. Nature Neuroscience 8, 1263–1268 (2005).

6. Wells, J., Kao, C., Mariappan, K., Albea, J., Jansen, E. D., Konrad, P. & Mahadevan-Jansen, A. Optical stimulation of neural tissue in vivo. Optics letters 30, 504–506 (2005).

7. Fekete, Z., Horváth, Á. C. & Zátonyi, A. Infrared neuromodulation: a neuroengineering perspective. Journal of Neural Engineering 17, 051003 (2020).

8. Jiang, S., Wu, X., Rommelfanger, N. J., Ou, Z. & Hong, G. Shedding light on neurons: optical approaches for neuromodulation. National Science Review (2022).

9. Bansal, A., Shikha, S. & Zhang, Y. Towards translational optogenetics. Nature Biomedical Engineering, 1-21 (2022).

10. Canales, A., Park, S., Kilias, A. & Anikeeva, P. Multifunctional fibers as tools for neuroscience and neuroengineering. Accounts of Chemical Research 51, 829–838 (2018).

11. Horváth, Á. C., Boros, Ö. C., Beleznai, S., Sepsi, Ö., Koppa, P. & Fekete, Z. A multimodal microtool for spatially controlled infrared neural stimulation in the deep brain tissue. Sensors and Actuators B: Chemical 263, 77–86 (2018).

12. Won, S. M., Cai, L., Gutruf, P. & Rogers, J. A. Wireless and battery-free technologies for neuroengineering. Nature Biomedical Engineering (2021).

13. Marin, C. & Fernández, E. Biocompatibility of intracortical microelectrodes: current status and future prospects. Frontiers in Neuroengineering 3, 8 (2010).

14. Park, S., Loke, G., Fink, Y. & Anikeeva, P. Flexible fiber-based optoelectronics for neural interfaces. Chemical Society Reviews 48, 1826–1852 (2019).

15. Bernstein, J. G., Garrity, P. A. & Boyden, E. S. Optogenetics and thermogenetics: technologies for controlling the activity of targeted cells within intact neural circuits. Current opinion in neurobiology 22, 61–71 (2012).

16. Wells, J., Kao, C., Konrad, P., Milner, T., Kim, J., Mahadevan-Jansen, A. & Jansen E. D. Biophysical mechanisms of transient optical stimulation of peripheral nerve. Biophysical Journal 93, 2567–2580 (2007).

17. Cayce, J. M., Wells, J. D., Malphrus, J. D., Kao, C., Thomsen, S., Tulipan, N. B., Konrad, P. E., Jansen, E. D. & Mahadevan-Jansen, A. Infrared neural stimulation of human spinal nerve roots in vivo. Neurophotonics, 2, 015007 (2015).

18. Wells, J. D., Kao, C., Jansen, E. D., Konrad, P. E. & Mahadevan-Jansen, A. Application of infrared light for in vivo neural stimulation. Journal of Biomedical Optics 10, 064003 (2005).

19. Xu, A. G., Qian, M., Tian, F., Xu, B., Friedman, R. M., Wang, J., Song, X., Sun, Y., Chernov, M. M., Cayce, J. M., Jansen, E. D., Mahadevan-Jansen, A., Zhang, X., Chen, G. & Roe, A. W. Focal infrared neural stimulation with high-field functional MRI: A rapid way to map mesoscale brain connectomes. Science Advances 5, eaau7046 (2019).

20. Jeong, J. W., McCall, J. G., Shin, G., Zhang, Y., Al-Hasani, R., Kim, M., Li, S., Sim, J. Y., Jang, K. I., Shi, Y., Hong, D. Y., Liu Y., Schmitz G. P., Xia L., He Z., Gamble P., Ray W. Z., Huang Y., Bruchas M. R. & Rogers J. A. Wireless optofluidic systems for programmable in vivo pharmacology and optogenetics. Cell 162, 662–674 (2015).

21. Antonini, M.J., Sahasrabudhe, A., Tabet, A., Schwalm, M., Rosenfeld, D., Garwood, I., Park, J., Loke, G., Khudiyev, T., Kanik, M., Corbin, N., Canales, A., Jasanoff, A., Fink, Y. & Anikeeva, P. Customizing MRI-Compatible Multifunctional Neural Interfaces through Fiber Drawing. Advanced Functional Materials 31, 2104857 (2021).

22. Zou, L., Tian, H., Guan, S., Ding, J., Gao, L., Wang, J. & Fang, Y. Self-assembled multifunctional neural probes for precise integration of optogenetics and electrophysiology. Nature Communications 12, 5871 (2021).

23. Kim, C. Y., Ku, M. J., Qazi, R., Nam, H. J., Park, J. W., Nam, K. S., Oh, S., Kang, I., Jang, J. H., Kim, W. Y., Kim, J. H. & Jeong, J. W. Soft subdermal implant capable of wireless battery charging and programmable controls for applications in optogenetics. Nature Communications 12, 535 (2021).

24. Shi, S., Xu, A. G., Rui, Y. Y., Zhang, X., Romanski, L. M., Gothard, K. M. & Roe, A. W. Infrared neural stimulation with 7T fMRI: A rapid in vivo method for mapping cortical connections of primate amygdala. NeuroImage 231, 117818 (2021).

25. Yoo, M., Koo, H., Kim, M., Kim, H. I. & Kim, S. Near-infrared stimulation on globus pallidus and subthalamus. Journal of Biomedical Optics 18, 128005 (2013).

26. Horváth, Á. C., Borbély, S., Boros, Ö. C., Komáromi, L., Koppa, P., Barthó, P., & Fekete, Z. Infrared neural stimulation and inhibition using an implantable silicon photonic microdevice. Microsystems & Nanoengineering 6, 44 (2020).

27. Abaya, T. V. F., Diwekar, M., Blair, S., Tathireddy, P., Rieth, L., Clark, G. A. & Solzbacher, F. Characterization of a 3D optrode array for infrared neural stimulation. Biomedical Optics Express 3, 2200–2219 (2012).

28. Woyessa, G., Nielsen, K., Stefani, A., Markos, C. and Bang, O. Temperature insensitive hysteresis free highly sensitive polymer optical fiber Bragg grating humidity sensor. Optics Express 24, 1206–1213 (2016).

29. Meneghetti, M., Petersen, C. R., Hansen, R. E., Adamu, A. I., Bang, O. & Markos, C. Thermally tunable dispersion modulation in a chalcogenide-based hybrid optical fiber. Optics Letters 46, 2533–2536 (2021).

30. Pranti, A.S., Schander, A., Bödecker, A. & Lang, W. PEDOT: PSS coating on gold microelectrodes with excellent stability and high charge injection capacity for chronic neural interfaces. Sensors and Actuators B: Chemical 275, 382–393 (2018).

31. Markos, C., Stefani, A., Nielsen, K., Rasmussen, H.K., Yuan, W. & Bang, O. High-T g TOPAS microstructured polymer optical fiber for fiber Bragg grating strain sensing at 110 degrees. Optics express 21, 4758–4765 (2013).

32. Dickinson, B. L., UDEL® Polysulfone for medical applications. Journal of Biomaterials Applications 3, 605–634 (1988).

33. Greer, A. I. M., Vasiev, I., Della-Rosa, B. & Gadegaard, N. Fluorinated ethylene– propylene: a complementary alternative to PDMS for nanoimprint stamps. Nanotechnology 27, 155301 (2016).

34. Agrawal, N. & Ugaz, V. M. A buoyancy-driven compact thermocycler for rapid PCR. Clinics in Laboratory Medicine 27, 215–223 (2007).

35. Sui, K., Meneghetti, M., Kaur, J., Sørensen, J.F., Berg, R. W. & Markos, C. Adaptive polymer fiber neural device for drug delivery and enlarged illumination angle for neuromodulation. Journal of Neural Engineering 19, 016035 (2022).

36. Henrie, J. A. & Shapley, R. LFP power spectra in V1 cortex: the graded effect of stimulus contrast. Journal of Neurophysiology 94, 479–490 (2005).

37. Englitz, B., David, S. V., Sorenson, M. D. & Shamma, S. A. MANTA—an open-source, high density electrophysiology recording suite for MATLAB. Frontiers in Neural Circuits 7, 69 (2013).

38. Cayce, J. M., Friedman, R. M., Chen, G., Jansen, E. D., Mahadevan-Jansen, A. & Roe, A.W. Infrared neural stimulation of primary visual cortex in non-human primates. Neuroimage 84, 181–190 (2014).

39. Priori, A., Foffani, G., Rossi, L. & Marceglia, S. Adaptive deep brain stimulation (aDBS) controlled by local field potential oscillations. Experimental Neurology 245, 77–86 (2013).

40. Averna, A., Marceglia, S., Arlotti, M., Locatelli, M., Rampini, P., Priori, A. & Bocci, T. Influence of inter-electrode distance on subthalamic nucleus local field potential recordings in Parkinson’s disease. Clinical Neurophysiology 133, 29–38 (2022).

41. Airaksinen, A. M., Niskanen, J. P., Chamberlain, R., Huttunen, J. K., Nissinen, J., Garwood, M., Pitkänen, A. & Gröhn, O. Simultaneous fMRI and local field potential measurements during epileptic seizures in medetomidine-sedated rats using raser pulse sequence. Magnetic Resonance in Medicine 64, 1191–1199 (2010).

42. Mao, X., Cao, T. & Li, A. Olfactory Receptors: Methods and Protocols Ch. 14 (Humana Press, New York, 2018).

43. Despouy, E., Curot, J., Reddy, L., Nowak, N., Deudon, M., Sol, J. C., Lotterie, J. A., Denuelle, M., Maziz, A., Bergaud, C., Thorpe, S. J., Valton, L. & Barbeau, E. J. Recording local field potential and neuronal activity with tetrodes in epileptic patients. Journal of Neuroscience Methods 341, 108759 (2020).

44. Itskov, V., Pastalkova, E., Mizuseki, K., Buzsaki, G. & Harris, K. D. Theta-mediated dynamics of spatial information in hippocampus. Journal of Neuroscience 28, 5959–5964 (2008).

45. Kropff, E., Carmichael, J. E., Moser, E. I. & Moser, M. B. Frequency of theta rhythm is controlled by acceleration, but not speed, in running rats. Neuron 109, 1029–1039 (2021).

46. Rogers, W. J. Sterilisation of Biomaterials and Medical Devices Ch. 7 (Woodhead Publishing, 2012)

47. Thermo Fisher Scientific, Teflon and other Fluorinated Ethylene Propylene (FEP) Labware, https://www.thermofisher.com/dk/en/home/life-science/lab-plasticware-supplies/plastic-material-selection/teflon-fep-labware.html

48. Mubarak, M. T., Ozsahin, I. & Ozsahin, D. U. Evaluation of sterilization methods for medical devices, paper presented in the 2019 Advances in Science and Engineering Technology International Conferences (ASET), Dubai, United Arab Emirates, 26 March-10 April 2019

49. Kozai, T. D. & Vazquez, A. L. Photoelectric artefact from optogenetics and imaging on microelectrodes and bioelectronics: new challenges and opportunities. Journal of Materials Chemistry B 3, 4965–4978 (2015).

50. Akrami, P., Adamu, A. I., Woyessa, G., Rasmussen, H. K., Bang, O. & Markos, C. All-polymer multimaterial optical fiber fabrication for high temperature applications. Optical Materials Express 11, 345–354 (2021).

51. El Waly, B., Escarrat, V., Perez-Sanchez, J., Kaur, J., Pelletier, F., Collazos-Castro, J. E. & Debarbieux, F. Intravital assessment of cells responses to conducting polymer-coated carbon microfibres for bridging spinal cord injury. Cells 10, 73 (2021)

52. Kaur, J. & Berg, R. W. Viral strategies for targeting spinal neuronal subtypes in adult wild-type rodents. Preprint at https://assets.researchsquare.com/files/rs-1050494/v1_covered.pdf (2021).

53. Kaur, J., Rauti, R. & Nistri, A. Nicotine-mediated neuroprotection of rat spinal networks against excitotoxicity. European Journal of Neuroscience 47, 1353–1374 (2018)

54. Kaur, J., Mazzone, G. L., Aquino, J. B. & Nistri, A. Nicotine neurotoxicity involves low Wnt1 signaling in spinal locomotor networks of the postnatal rodent spinal cord. International Journal of Molecular Sciences 22, 9572 (2021).

55. Wieliczka, D. M., Weng, S. & Querry, M. R. Wedge shaped cell for highly absorbent liquids: infrared optical constants of water. Applied Optics 28, 1714–1719 (1989).

56. Paltauf, G. & Dyer, P. E. Photomechanical processes and effects in ablation. Chemical reviews 103, 487–518 (2003).

57. Ma, W., Liu, W. & Li, M. Analytical heat transfer model for targeted brain hypothermia. International Journal of Thermal Sciences 100, 66–74 (2016).

58. Azhari, H. Basics of biomedical ultrasound for engineers App. A (John Wiley & Sons, New Jersey, 2010)

59. Thompson, A.C., Wade, S.A., Brown, W.G. & Stoddart, P.R. Modeling of light absorption in tissue during infrared neural stimulation. Journal of biomedical optics 17, 075002 (2012).

