## Supplementary Information for "Opsin-free optical neuromodulation and electrophysiology enabled by a soft monolithic infrared multifunctional neural interface"

**Supplementary Text**

1: Data post-processing

A notch filter (50 Hz) has been digitally applied to all electrophysiological recordings presented in the main text. Additionally, to highlight the neural activity "buried" in noise for the stimulations performed with 10, 12.5 and 15 mW optical power presented in Fig. 2, a low-pass filter with a cut-off frequency of 40 Hz has been applied prior to plotting. The lower signal-to-noise ratio in these recordings has been attributed to a combination of suboptimal contact between the electrodes and the neurons and the presence of multi-unit activity in these recording, which could be suppressed and distorted by the 300 Hz high-pass filter integrated in the recording setup used. The full recordings and examples of spikes are presented in Fig. S3-S6, respectively, both as recorded and after filtering.

2: Thermal and stress confinement conditions

Photomechanical effects in laser-tissue interaction are enhanced when thermal and stress confinement conditions are met, i.e. when the laser pulse width $t_p$ is smaller than the thermal and acoustic relaxation times $t_{th}$ and $t_{ac}$ of the tissue. These relaxation times can be calculated as [56]:

$$t_{th} = \frac{d^2}{4\chi}$$

and

$$t_{ac} = \frac{d}{c_s}$$

Where $d$ is the smallest dimension of the irradiated volume, $\chi$ is the tissue thermal diffusivity coefficient (0.129 mm$^2$/s in brain tissue [57]) and $c_s$ is the speed of sound (1550 m/s in brain tissue [58]). For the experiments described in this manuscript, where the light delivered to the brain has a relatively broad spectrum (1800-2100 nm), $d$ cannot be directly defined. For wavelengths that are less absorbed from the brain (i.e. absorption coefficient $\alpha < 95$ cm$^{-1}$ for water [59]), $d$ is equal to the fiber core diameter (i.e. 105 µm). For wavelengths with $\alpha > 95$ cm$^{-1}$, which constitute ~30% of the delivered light, $d$ is instead equal to the penetration depth of light in the tissue (i.e. the distance at which the intensity of light drops to 1/e). As the maximum value of $\alpha$ for water in the 1800-2100 nm band is equal to 130.6 cm$^{-1}$ [55], it is however possible to determine that $d$ is strictly comprised between 76 and 105 µm. The corresponding ranges for the relaxation times are ~10-100 ms for $t_{th}$ and ~45-70 ns for $t_{ac}$. The supercontinuum laser, with $t_p$ being picosecond range, therefore meets the conditions for both confinement regimes (thermal and stress).

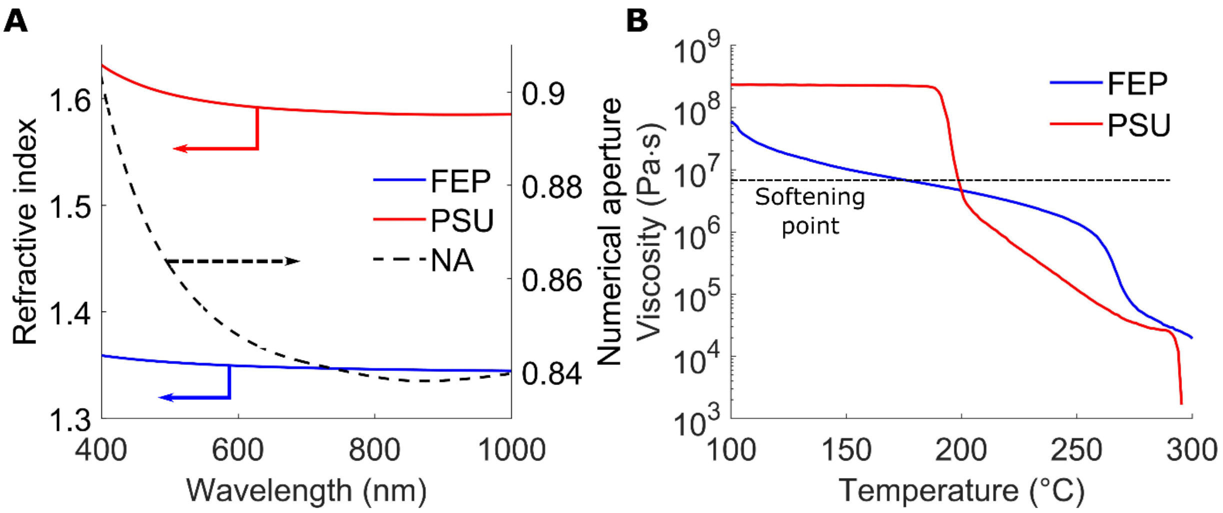


**Fig. S1.**

(**A**) Viscosity profile of the two polymers used to develop the SPOF. The dashed line indicates the Littleton Softening Point ($10^{6.6}$ Pa·s). (**B**) Refractive indexes (solid lines) of the two polymers used to develop the SPOF and calculated NA of the SPOF.

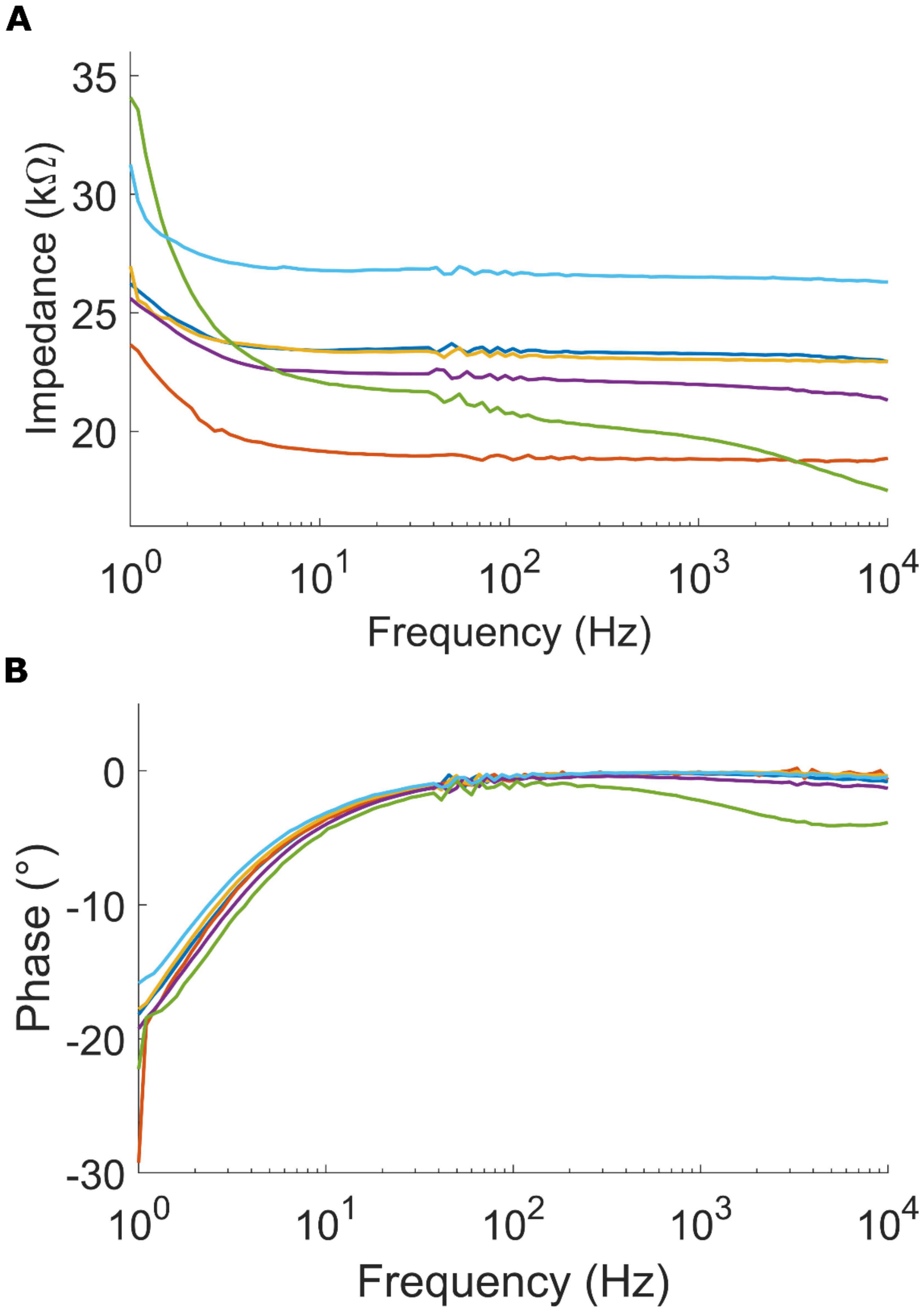


**Fig. S2.**

(**A**) Impedance and (**B**) phase recordings for 6 electrodes integrated into the developed neural interfaces.

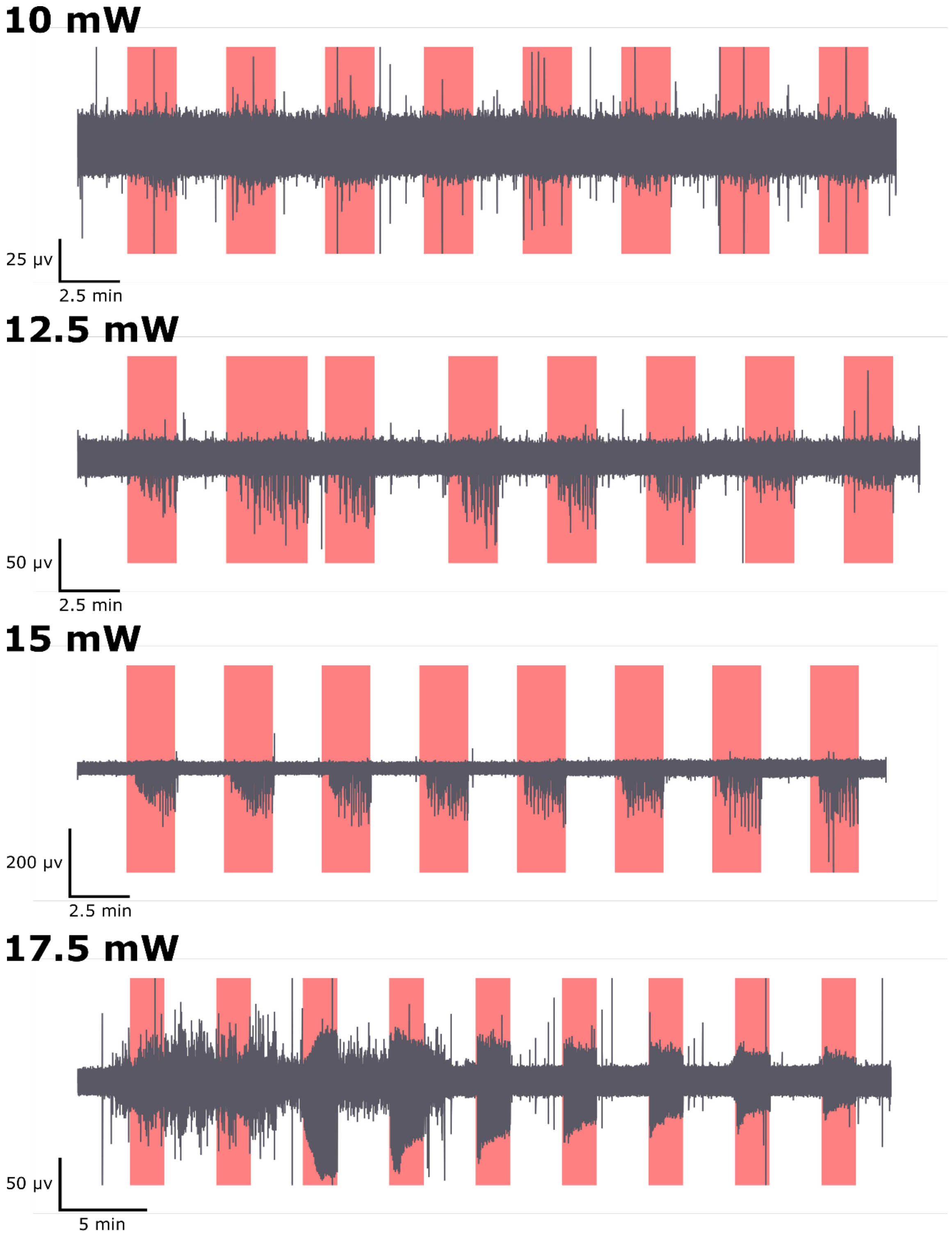


**Fig. S3.**
Electrophysiological recordings during several stimulation cycles (red regions), as recorded.

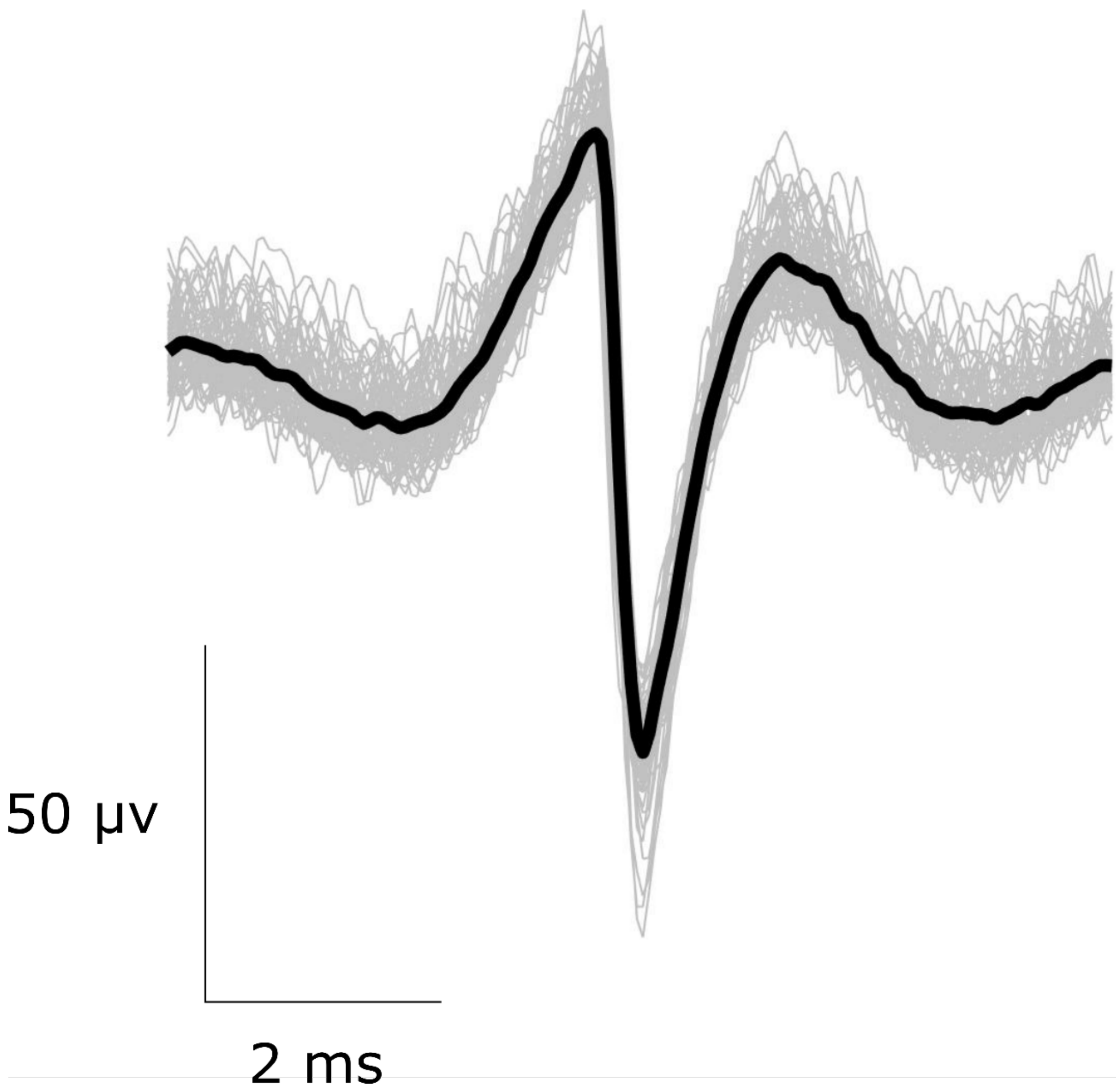


**Fig. S4.**

Average spike activity recording during the 17.5mW stimulation cycle. Light grey is the spikes' overlap, and black represents the average value, depicted in Fig. 2C (bottom) (n=62).

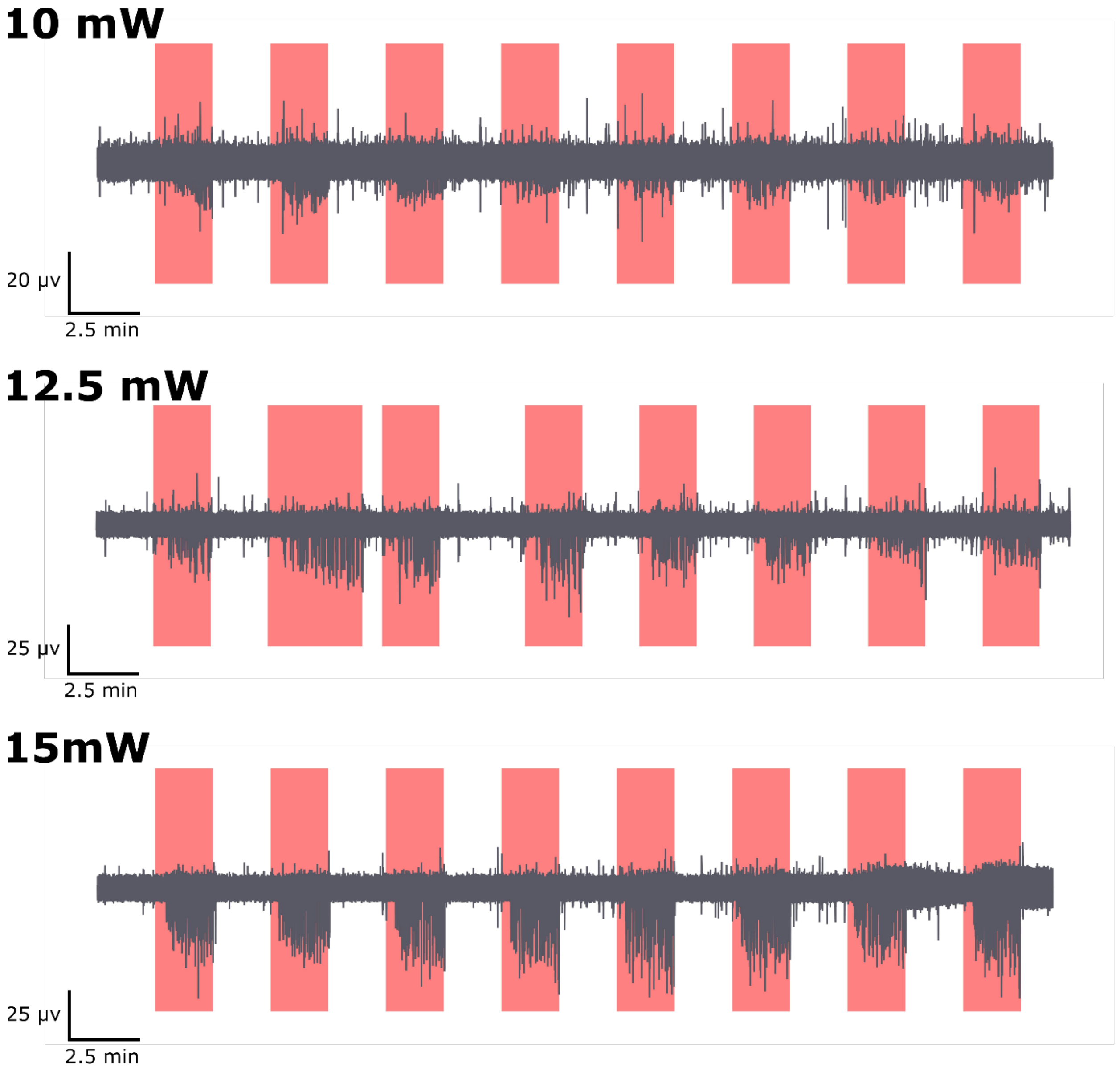


**Fig. S5.**
Electrophysiological recordings during several stimulation cycles (red regions) after applying a 40 Hz low-pass filter.

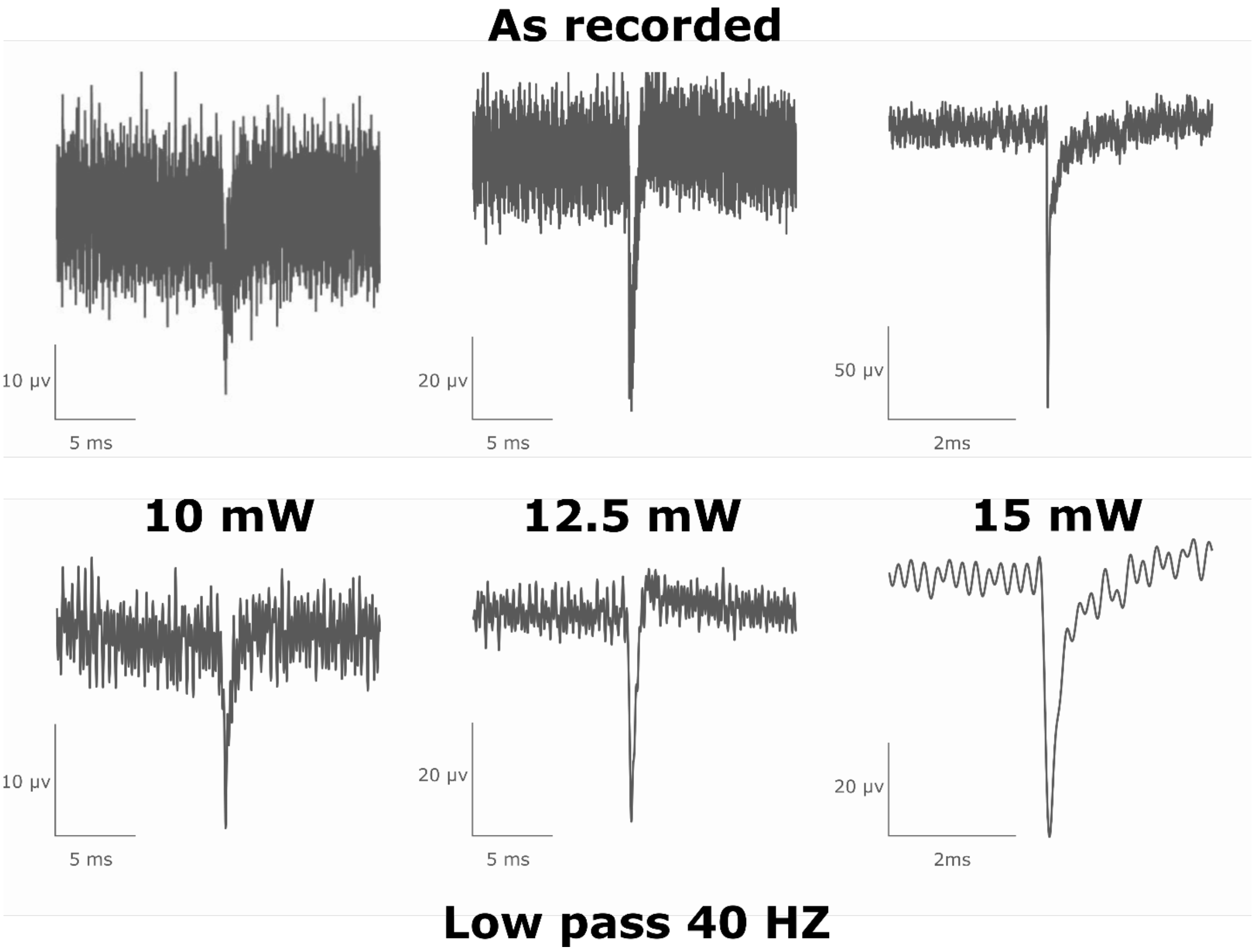


**Fig. S6.**
Example of spikes extracted by the recordings in Figure S5, both as recorded and after applying a digital low-pass filter to increase the signal-to-noise ratio.

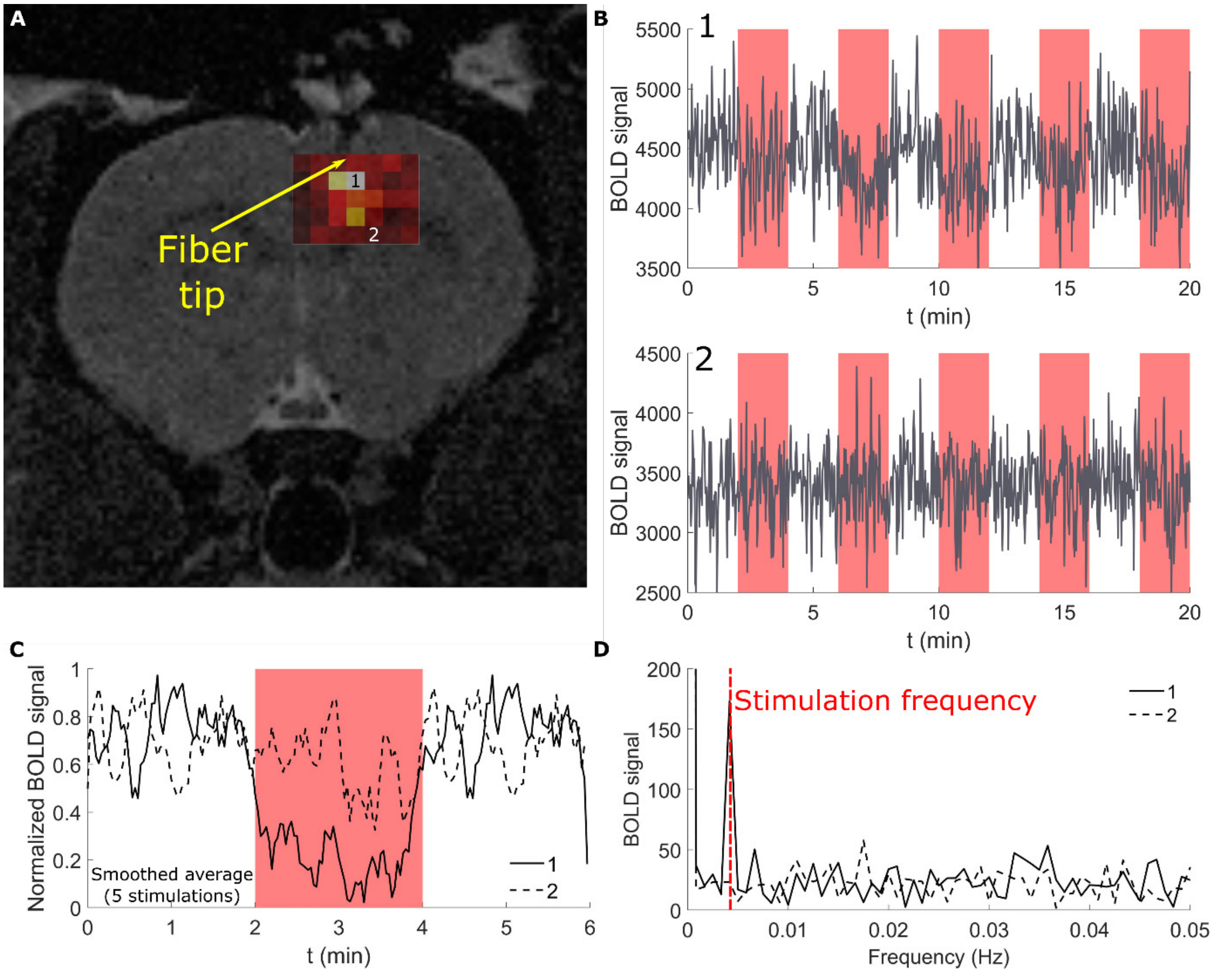


**Fig. S7.**

(**A**) Overlap between a structural MRI image and the signal intensity at the stimulation frequency obtained by fast Fourier transform of the time profile of an EPI scan (stimulation power: 15 mW). (**B**) Temporal profile of the BOLD signal during the scan for voxels 1 and 2 are indicated in (**A**) (stimulations in red). (**C**) Comparison of the smoothed average of BOLD signal over 5 stimulations between voxel 1 and voxel 2. (**D)** Comparison of the spectral components of BOLD signal between voxel 1 and voxel 2.

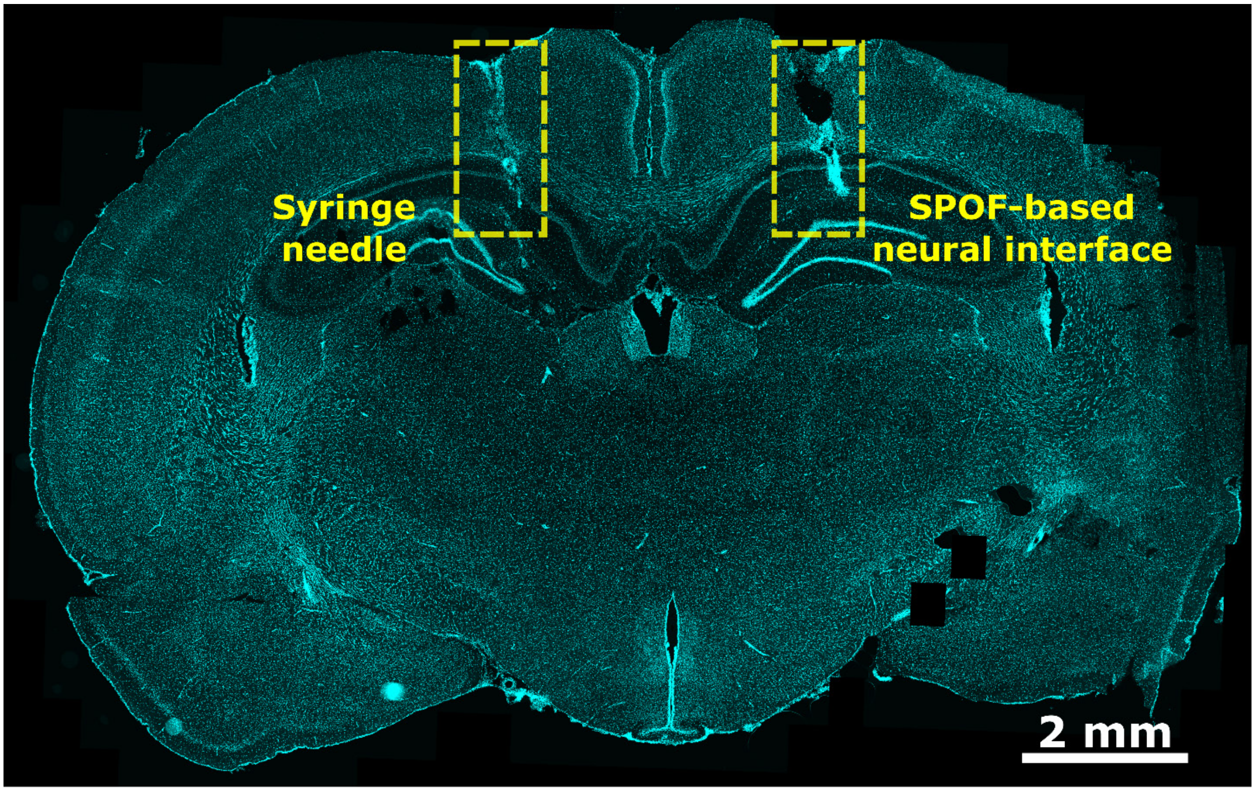


**Fig. S8.**
Brain slice microscope image (DAPI staining) showing the footprints after 4 weeks of implantation of a steel needle with tungsten µwires (left) and the SPOF-based neural interface (right).

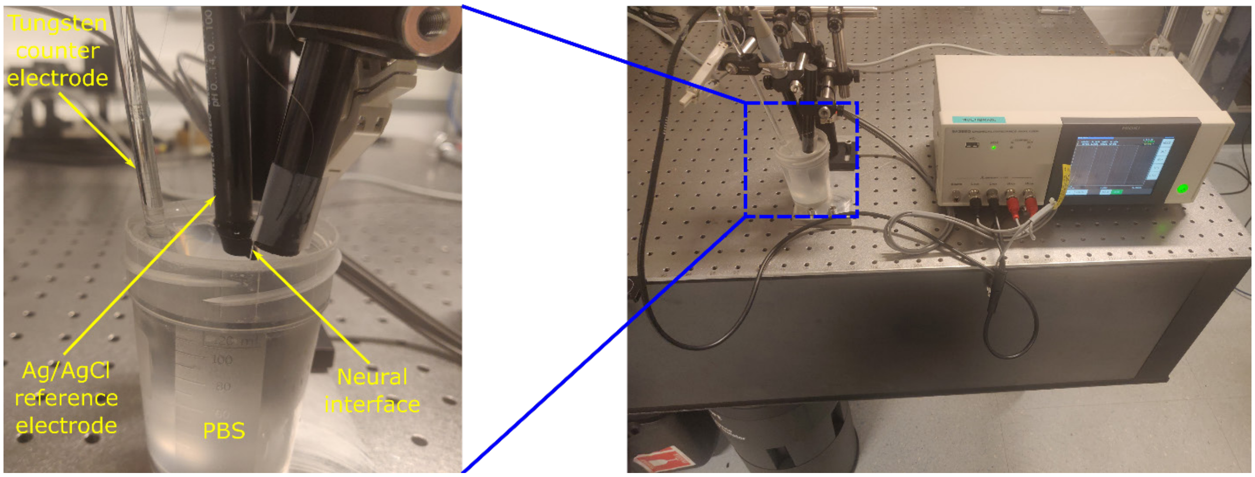


**Fig. S9.**
Setup used for electrochemical impedance spectroscopy measurements.

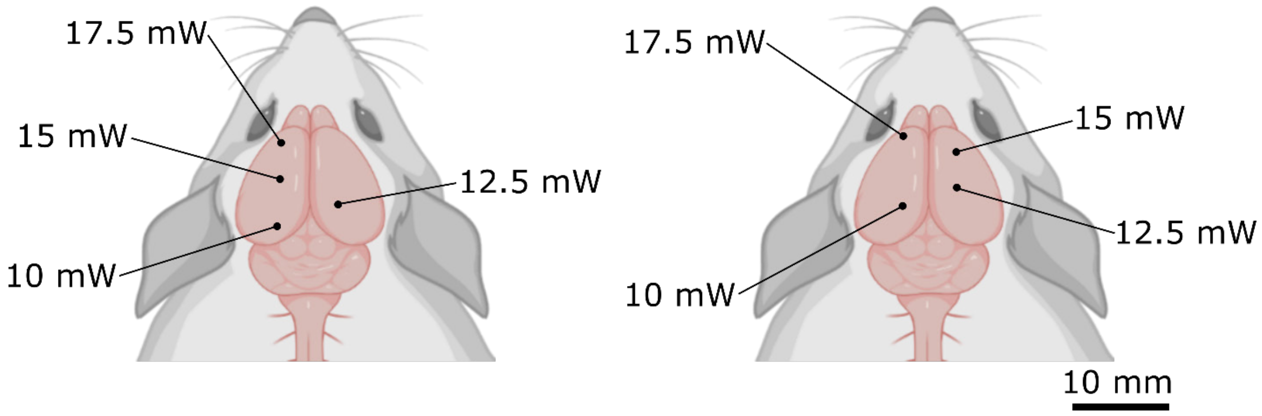


**Fig. S10.**
Locations in the cortex which were used for the two simultaneous INM and EE experiments (insertion depth of 1 mm for all locations).
